# Characterization of a novel intratracheal aerosol challenge model of *Brucella melitensis* in guinea pigs

**DOI:** 10.1101/457184

**Authors:** M.E. Hensel, D.G. Garcia-Gonzalez, S.P. Chaki, J. Samuel, A.M. Arenas-Gamboa

## Abstract

*B. melitensis* is considered the most virulent of the *Brucella* species, and a need exists for an improved laboratory animal model of infection that mimics natural transmission and disease. Guinea pigs are highly susceptible to infection with *Brucella* spp. and develop a disease syndrome that mimics natural disease after aerosol inoculation. Intratracheal inoculation is a targeted means of generating aerosols that offer advantages over aerosol chamber delivery. To establish this delivery method, female, Hartley guinea pigs were infected via intratracheal inoculation with PBS or 16M *B. melitensis* at low dose (10^1^ to 10^3^) or high dose (10^6^ to 10^8^) and monitored for 30 days for signs of disease. Guinea pigs in the high dose groups developed fever between 12-17 days post-inoculation. Bacteria were recovered from the spleen, liver, lymph nodes, lung, and uterus at 30-days post-inoculation and demonstrated dose dependent mean increases in colonization and pathologic changes consistent with human brucellosis. To study the kinetics of extrapulmonary dissemination, guinea pigs were inoculated with 10^7^ CFU and euthanized at 2-hours post inoculation and at weekly intervals for 3 weeks. 5.8×10^5^ to 4.2×10^6^ CFU were recovered from the lung 2 hours post-inoculation indicating intratracheal inoculation is an efficient means of infecting guinea pigs. Starting at 1-week post inoculation bacteria were recovered from the aforementioned organs with time dependent mean increases in colonization. This data demonstrates that guinea pigs develop a disease syndrome that models the human manifestation of brucellosis, which makes the guinea pig a valuable model for pathogenesis studies.

**Author summary:** Brucellosis is caused by a gram-negative, intracellular bacterial pathogen with a worldwide distribution and affects up to half a million people per year. It is a neglected zoonosis that impacts not only animal welfare, but also exert economic pressure on afflicted individuals through loss of wages and decreased productivity. In people, recurrent fever, malaise, and anorexia accompanied by enlargement of the spleen and lymph nodes are common clinical symptoms of infection. The mouse model has been used extensively to study the pathogenesis of brucellosis, but there are drawbacks to extrapolating studies in mice to develop vaccines or therapeutics for people. Mice are frequently inoculated via intraperitoneal injection, which is an artificial means of producing disease that does not mimic natural transmission or disease features, such as fever. An animal model is needed that can be infected through natural transmission routes and subsequently develop a syndrome that matches clinical disease seen in people in order to study the pathogenesis of disease and to develop vaccines and therapeutics. The guinea pig offers an improvement on the mouse model because it can be infected via aerosol inoculation and develops fever, a humoral immune response, systemic colonization, and macroscopic and microscopic lesions of disease. As such, guinea pigs could be used a more biologically relevant model for evaluation of host-pathogen interactions.

## Introduction

Brucellosis is a disease caused by a gram-negative coccobacillus of the genus *Brucella* and is a zoonotic pathogen that has a worldwide distribution [1]. Of the twelve currently recognized *Brucella* species, *Brucella melitensis* is considered the most virulent [2]. The natural hosts of *B. melitensis* are sheep and goats [2]. The primary clinical presentation in affected small ruminants are abortion, stillbirths, and decreased fertility; bacteria are shed in large numbers after abortions in the placenta or through secretory products like milk [2]. People are commonly exposed through aerosols or by ingestion of unpasteurized milk or milk products [2]. In humans, clinical brucellosis typically manifests as relapsing periods of fever, malaise, and inappetance [2]. More severe complications such as disease of the reproductive, osteoarticular, cardiovascular, or nervous systems are also possible [2, 3].

Aerosols are a common means of transmission in people and animals and inhalation of bacteria leads to colonization of the reticuloendothelial organs such as the spleen, liver, and lymph nodes [2]. Certain occupations are at a greater risk of exposure due to close proximity with animals including veterinarians, farmers, and abattoir workers [2]. Humans who are exposed to aerosols generated following an animal abortion event are often exposed to up to 10^9^ colony forming units (CFU), but a dose of 10-100 CFU is reported to generate disease [2, 4]. Due to the ease of aerosolization and the low infectious dose, *B. melitensis* could potentially be weaponized and is designated a Category B agent by the Centers for Disease Control and Prevention [4].

Animal models utilized to study human brucellosis include mice, guinea pigs, rabbits, rats, and nonhuman primates [5]. Mice are currently the most commonly used model for brucellosis due to the ready availability of many genetic and immunologic tools [5]. A drawback to murine research is the large number of infectious organisms required to induce disease, which is well above the dose required to cause infection in people, and mice do not develop fever [6, 7]. Additionally, the most common means of inoculating mice with *Brucella* is intraperitoneal injection, which is not a means of natural transmission and thus the results of these experiments may not be as relevant. Guinea pigs were the animal model of choice to study the pathogenicity of *Brucella* species from the early 1900s to 1960 but were supplanted by the mouse model [8-10]. Similar to mice, guinea pigs can be infected by a variety of routes including intraperitoneal, intramuscular, subcutaneous, and inhalation. In contrast to mice, guinea pigs not only develop systemic disease but also demonstrate clinical signs of infection that include fever [11]. A need exists for an animal model that can be infected via aerosol transmission and replicate key features of human disease.

The experiments described herein represent a novel approach to understand the pathogenesis of aerosol transmission in a guinea pig model including the dose response to infection, kinetics of dissemination after aerosol exposure, and macroscopic and microscopic pathologic findings. Previous studies have indicated that guinea pigs are a physiologically relevant model and with an updated approach to inoculation, the guinea pig could be used to investigate host-pathogen interactions.

## Results

### Passage through the MicroSprayer^®^ does not adversely affect bacterial viability

This study utilized the PennCentury^TM^ MicroSprayer because it is a targeted means of generating aerosols and has been used successfully to inoculate mice with bacterial pathogens [12]. The MicroSprayer^®^ device has not been previously used to inoculate guinea pigs. Our first objective was to determine if passage of the inoculum through the MicroSprayer^®^ affected the bacterial viability. Bacterial suspensions of each dose were sprayed through the device and collected into 900 µl of PBS, serially diluted, and plated on TSA to calculate the number of viable bacteria. Bacterial viability was minimally affected by passage through the device. As an example, the original inoculum for guinea pigs in the 10^7^ group contained 4.4×10^7^ CFU/50 µl and after passage through the MicroSprayer^®^ 4.1×10^7^ CFU/50 µl were recovered (Table 1). This study proves that the device is a reliable means of generating an infectious aerosol and passage through the MicroSprayer^®^ does not adversely affect the viability of the bacteria.

**Table 1.** Bacterial viability following passage through the MicroSprayer^®^ Aerosolizer.

| Dose group | Original | MicroSprayer® |
| --- | --- | --- |
| 10<sup>1</sup> | 2.0x10<sup>2</sup> | 1x10<sup>2</sup> |
| 10<sup>2</sup> | 2.3x10<sup>3</sup> | 2.0x10<sup>2</sup> |
| 10<sup>3</sup> | 7.9x10<sup>3</sup> | 1.65x10<sup>3</sup> |
| 10<sup>6</sup> | 4.40x10<sup>6</sup> | 3.80x10<sup>6</sup> |
| 10<sup>7</sup> | 4.40x10<sup>7</sup> | 4.10x10<sup>7</sup> |
| 10<sup>8</sup> | 4.80x10<sup>8</sup> | 4.10x10<sup>8</sup> |

### Intratracheal inoculation with 16M *Brucella melitensis* results in systemic disease

Having established that the MicroSprayer^®^ does not adversely affect bacterial viability, we next evaluated the ability of the device to inoculate guinea pigs with low doses (10^1^, 10^2^,10^3^) or high doses (10^6^, 10^7^, 10^8^) of *B. melitensis* 16M. After intratracheal inoculation with *B. melitensis*, guinea pigs were monitored for signs of clinical disease including fever, loss of appetite, respiratory disease (ocular discharge, increased respiratory effort), and lethargy. Brucellosis is a disease of high morbidity but low mortality and, as expected, intratracheal inoculation did not result in any deaths in any dose group despite evidence of systemic infection. However, guinea pigs in the 10^8^ group had more severe clinical signs including roughened hair coat, ocular discharge, and lethargy. Body weight was not affected by infection in any dose group, and all guinea pigs continued to gain weight throughout the study period (data not shown). Guinea pigs inoculated with PBS or the low doses (10^1^, 10^2^,10^3^) of *B. melitensis* did not develop fever or other clinical signs of brucellosis at any time point (data not shown). In the high dose groups, the onset of fever (temperature ≥39.5°C) developed in a dose-dependent manner beginning at day 16 post-infection (Fig. 1). Approximately 75% of the animals in the 10^6^ and 10^7^ groups developed fever with an undulant pattern. Based on the kinetics study, the earliest onset of fever appears to be 12-days post-inoculation (data not shown). The average daily temperature was significantly increased (*P* <0.05) in the 10^6^ and 10^7^ groups between days 16 to 24 compared to the uninfected control group. The guinea pigs in the 10^8^ group did not develop fever to the same level but 2 animals had a single episode of fever. We ascribe the lack of fever response in the 10^8^ group to overwhelming disease that resulted in sepsis.

**Figure 1.**
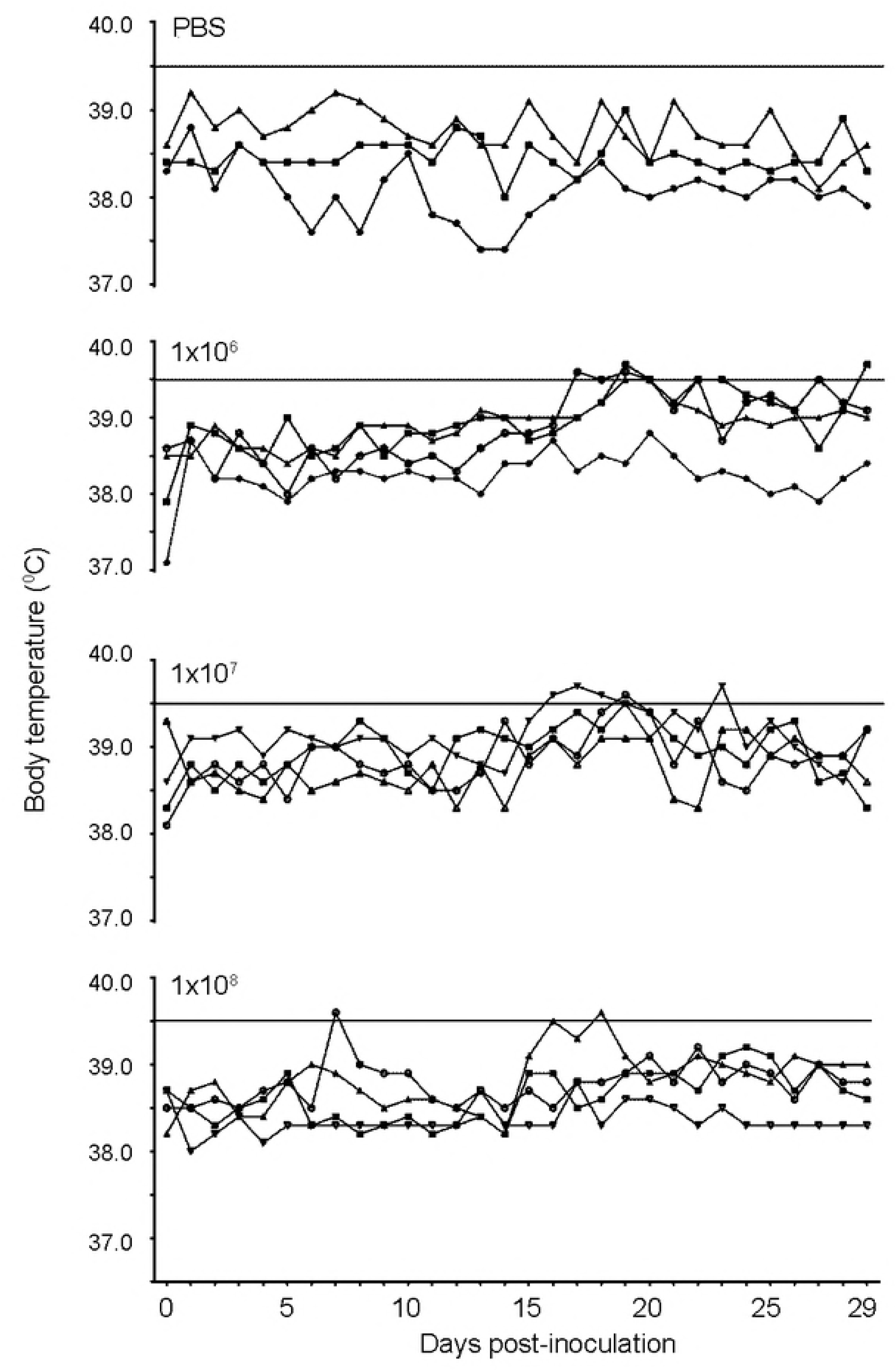
Body temperature changes in guinea pigs after intratracheal inoculation PBS, 10^6^, 10^7^, and 10^8^ *B. melitensis* 16M. Guinea pigs were inoculated using the MicroSprayer^®^ Aerosolizer with low dose (10^1^, 10^2^,10^3^), high dose (10^6^, 10^7^, 10^8^), or control (PBS) groups. Body temperature was monitored daily using an implantable subcutaneous IPTT-300 microchip. The solid line at 39.5° C indicates the threshold for fever. Guinea pigs in the 10^6^ and 10^7^ groups developed fever with an undulant pattern.

A hallmark of brucellosis in natural hosts and humans is splenomegaly. Previous aerosol studies with guinea pigs demonstrated the development of splenomegaly after infection [13]. In response to infection, spleen weight was significantly increased (p<0.0001) in the high dose group (10^6^, 10^7^, 10^8^) compared to the uninfected controls (Fig. 2A). The average spleen weight in the 10^6^, 10^7^, and 10^8^ group was 3.45 g, 2.96 g, and 3.33 g, respectively compared to 0.6 g in the control group. Spleen weight continuously increased over a four-week course of infection (Fig. 2B). While the liver is a frequent target of *B. melitensis*, infection is not associated with hepatomegaly in humans [14]. Similarly, the liver weight was not significantly different between dose groups or time points in guinea pigs (data not shown).

**Figure 2.**
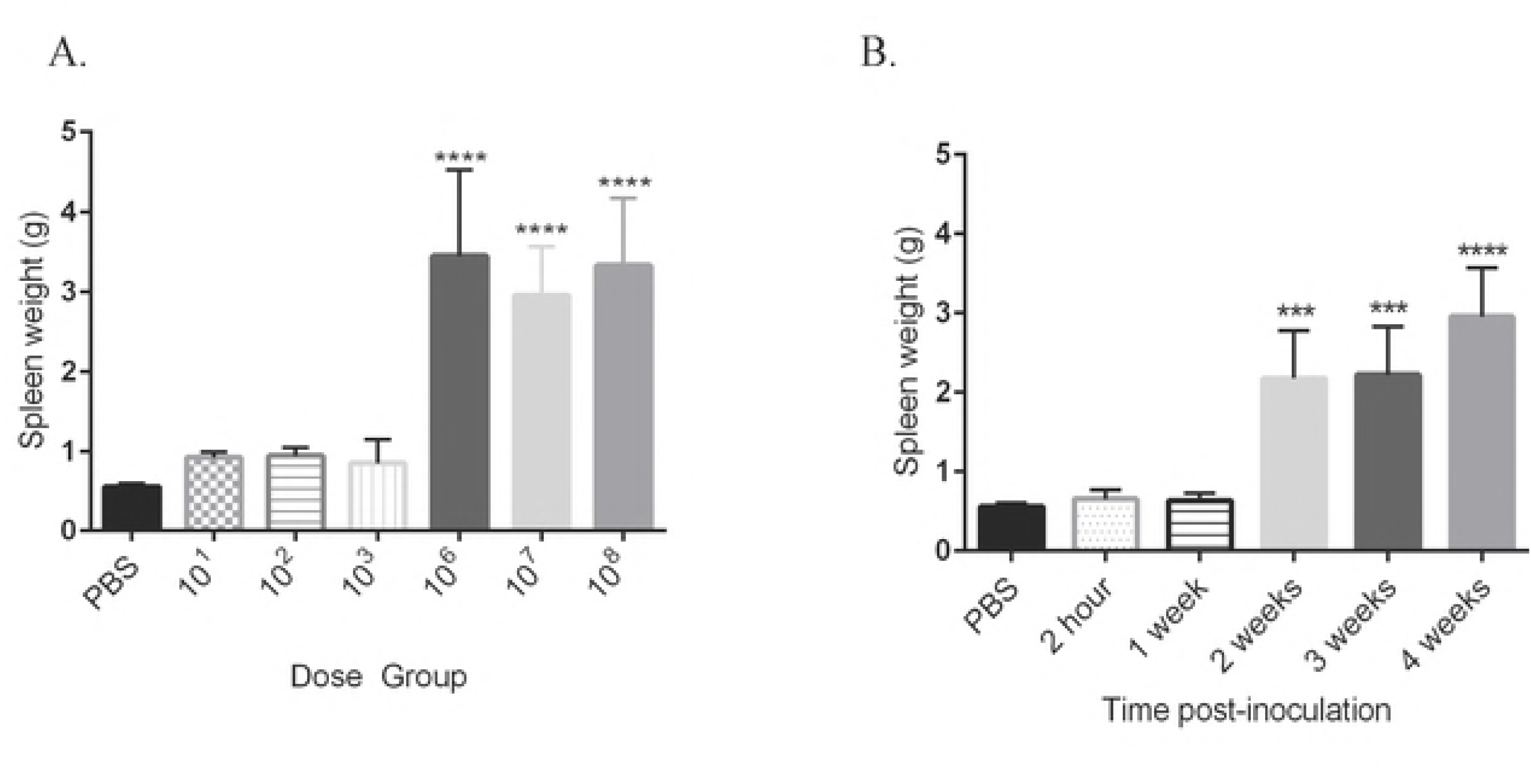
Splenic weights in guinea pigs inoculated with *B. melitensis* 16M or PBS. Splenomegaly was induced by high doses (10^6^, 10^7^, 10^8^) of *B. melitensis* by 30-days post-inoculation (A). Splenomegaly was detected as early as 2-weeks post-inoculation and increased through the study period (B). Data bars represent the mean spleen weight plus the standard deviation for all guinea pigs in each dose group. Mean spleen weight from each dose group (n=4) or time point (n=4) was compared to mean spleen weight of the uninfected control guinea pigs (n=3) and statistical significance was determined by ANOVA followed by Dunnett’s multiple-comparison test. Three asterisks, *P* <0.001. Four asterisks, *P* <0.0001.

### Guinea pigs infected with *B. melitensis* develop macroscopic and microscopic lesions

*Brucella* spp. have a tropism for tissues of the reticuloendothelial system and reproductive tract. To determine colonization following intratracheal inoculation, the spleen, liver, lung, cervical lymph node (CLN), tracheobronchial lymph node (TBLN) and uterus were collected for culture. Guinea pigs inoculated with either PBS or 10^1^ and 10^2^ CFU doses of *B. melitensis* did not result in colonization of any tissue examined. Animals in the 10^3^ and high dose groups (10^6^, 10^7^, 10^8^) demonstrated dose-dependent mean increases in CFU recovered per gram of the spleen, liver, lung, cervical lymph node, tracheobronchial lymph node, and uterus at 30- days post-inoculation (Fig. 3A-F). Following intratracheal inoculation, bacteria are rapidly disseminated to the spleen, draining lymph nodes, and uterus within 2-hours post-inoculation and could be recovered from the lung, CLN, and TBLN in 100% of the animals (Fig. 4C-F). The inoculum was evenly distributed throughout all lung lobes indicating that intratracheal inoculation generates a particle size that is able to reach the terminal airways (Fig. S1). Peak replication occurred at 3-weeks post-inoculation in the spleen, liver, and uterus (Fig. 4A,B,D). Replication continued to increase in the CLN and TBLN for the entire study period (Fig. 4E-F).

**Figure 3.**
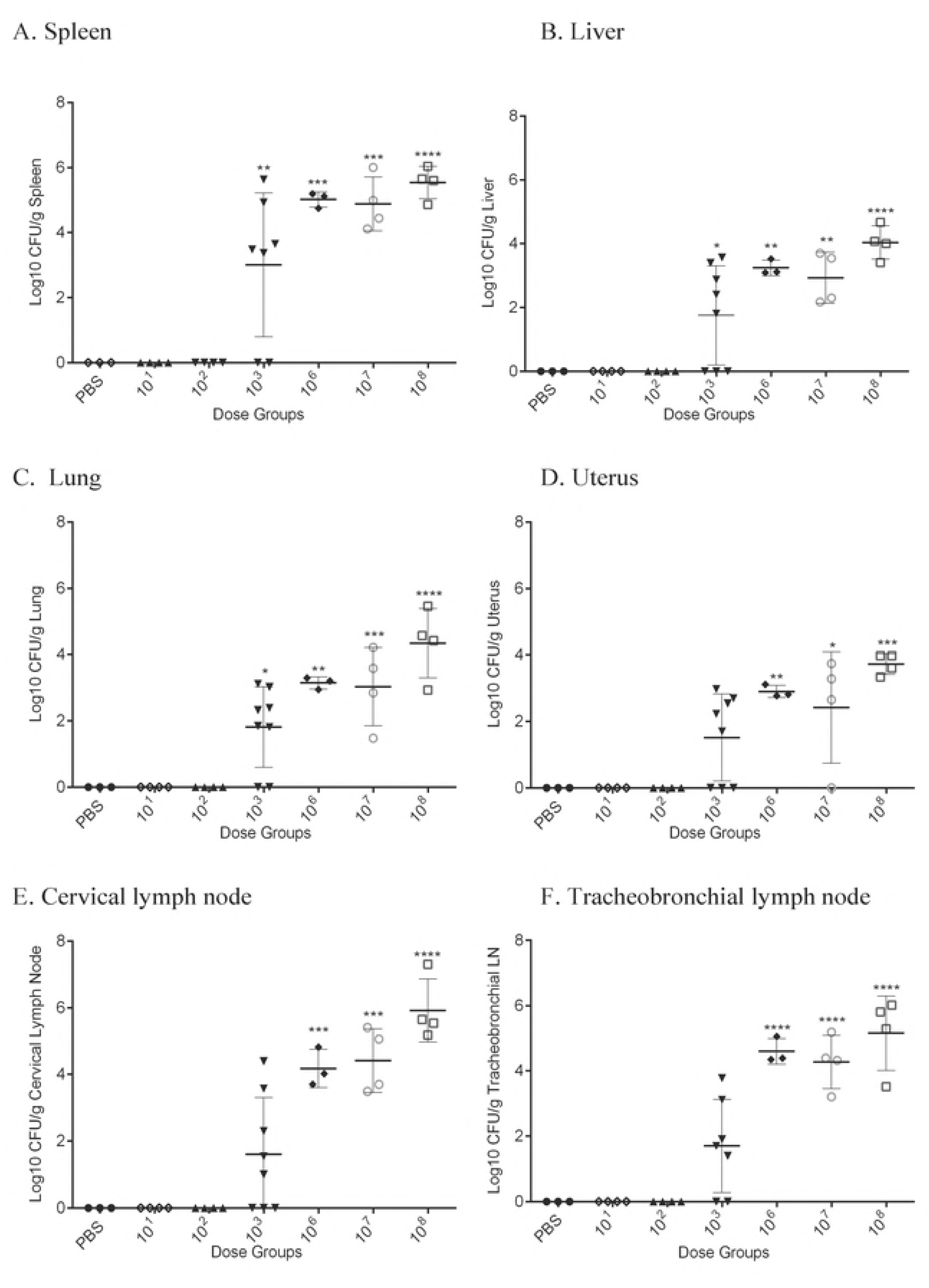
Intratracheal inoculation with *B. melitensis* 16M in female Hartley guinea pigs results in systemic infection. Guinea pigs were divided in 7 groups (n=4) consisting of low dose (10^1^, 10^2^,10^3^), high dose (10^6^, 10^7^, 10^8^), or control (PBS) groups. Guinea pigs were inoculated using the MicroSprayer^®^ Aerosolizer and were euthanized 30-days post-inoculation. Colonization was evaluated in the spleen (A), liver (B), lung (C), uterus (D), cervical lymph node (E), and tracheobronchial lymph node (F). The recovery of organisms is plotted as the total CFU/g (means ± standard deviation). Mean recovery per gram of tissue was compared between dose groups and uninfected control guinea pigs. Statistical significance was determined by ANOVA followed by Dunnett’s multiple comparisons. One asterisk, *P* < 0.05. Two asterisks, *P* <0.01. Three asterisks, *P* <0.001. Four asterisks, *P* <0.0001.

**Figure 4.**
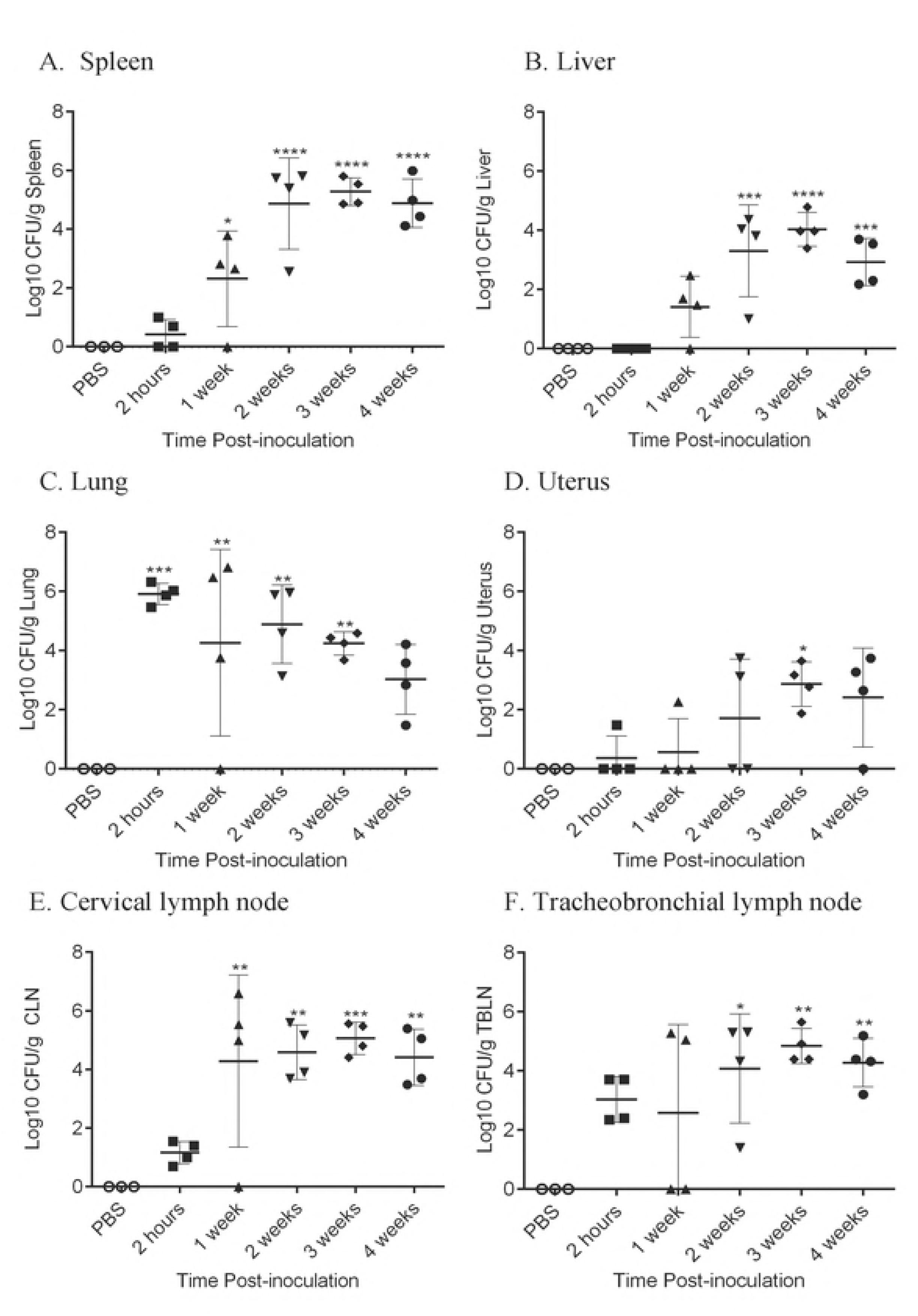
Kinetics of systemic infection of *B. melitensis* 16M in guinea pigs infected via intratracheal inoculation. Four female Hartley guinea pigs per time point group were inoculated intratracheally with 1×10^7^ CFU/50 µl. The initial lung colonization was evaluated 2-hours post-inoculation to determine the inhaled dose. Guinea pigs (n=4) were euthanized at 1,2,3, and 4-weeks post-inoculation to determine the numbers of *B. melitensis* in the spleen (A), liver (B), lung (C), uterus (D), cervical lymph node (E), and tracheobronchial lymph node (F). Mean recovery per gram of tissue was compared between time points and uninfected control guinea pigs. Statistical significance was determined by ANOVA followed by Dunnett’s multiple comparisons. One asterisk, *P* < 0.05. Two asterisks, *P* <0.01. Three asterisks, *P* <0.001. Four asterisks, *P* <0.0001.

The earliest gross lesions developed 2-weeks post-inoculation and included nodular lymphoid hyperplasia in the spleen, perinodal hemorrhage around the CLN, multifocal random 1-2 mm pale foci in the liver, and consolidation of the cranioventral lung lobes with multifocal 1-3 mm depressed gray foci scattered throughout the pulmonary parenchyma. A single animal in the 10^7^ group had a splenic abscess. No gross or microscopic lesions consistent with brucellosis were observed in any organ in the PBS control, 10^1^, or 10^2^ groups or at 2-hours post-inoculation. A grading system was developed to assess microscopic findings in the spleen, liver, lung, and uterus (Table S1). Application of the grading system demonstrated a significant increase (*P*<0.0001) in lesion severity based on average histologic score as the dose increased between the uninfected controls and high dose groups.

Sections were graded by a board-certified veterinary pathologist (MH). Lesions in all organs in the high dose groups (10^6^, 10^7^, 10^8^) increased in number, size, and severity by 30-days post-inoculation in a dose dependent manner. Histologic evaluation of the spleen revealed an inflammatory infiltrate of predominantly epithelioid macrophages with fewer neutrophils that effaced the normal architecture (Fig. 5). Similarly, the earliest lesion at 1-week post-inoculation were small foci of epithelioid macrophages in the red pulp that increased in size and number at 2 and 3-weeks post-inoculation. The cortex and medulla of the lymph node were also expanded by a large number of epithelioid macrophages (data not shown). The liver lesion was characterized by variably sized random foci of liquefactive and coagulative necrosis surrounded by neutrophilic and histiocytic inflammation and multifocal random microgranulomas composed of accumulations of histiocytes (Fig. 6). Portal areas were expanded by lymphocytes and plasma cells. The range of morphologic diagnoses seen in the guinea pigs is similar to those described in the liver of people infected with *B. melitensis* including lymphocytic portal hepatitis and microgranulomas. In addition, guinea pigs had foci of necrosis surrounded by macrophages and neutrophils, which may be similar to the noncaseating granulomas described by Young [15]. The earliest lesion in the lung at 1-week post-inoculation included expansion of the bronchus-associated lymphoid tissue (BALT), congestion of the alveolar walls, and edema. By 2-weeks post-inoculation, alveolar walls were thickened by an inflammatory infiltrate of macrophages and neutrophils surrounded by lymphocytes and plasma cells. At 3 to 4-weeks, the inflammatory infiltrate had coalesced into variably sized nodules of histiocytic and neutrophilic inflammation (Fig. S2).

**Figure 5.**
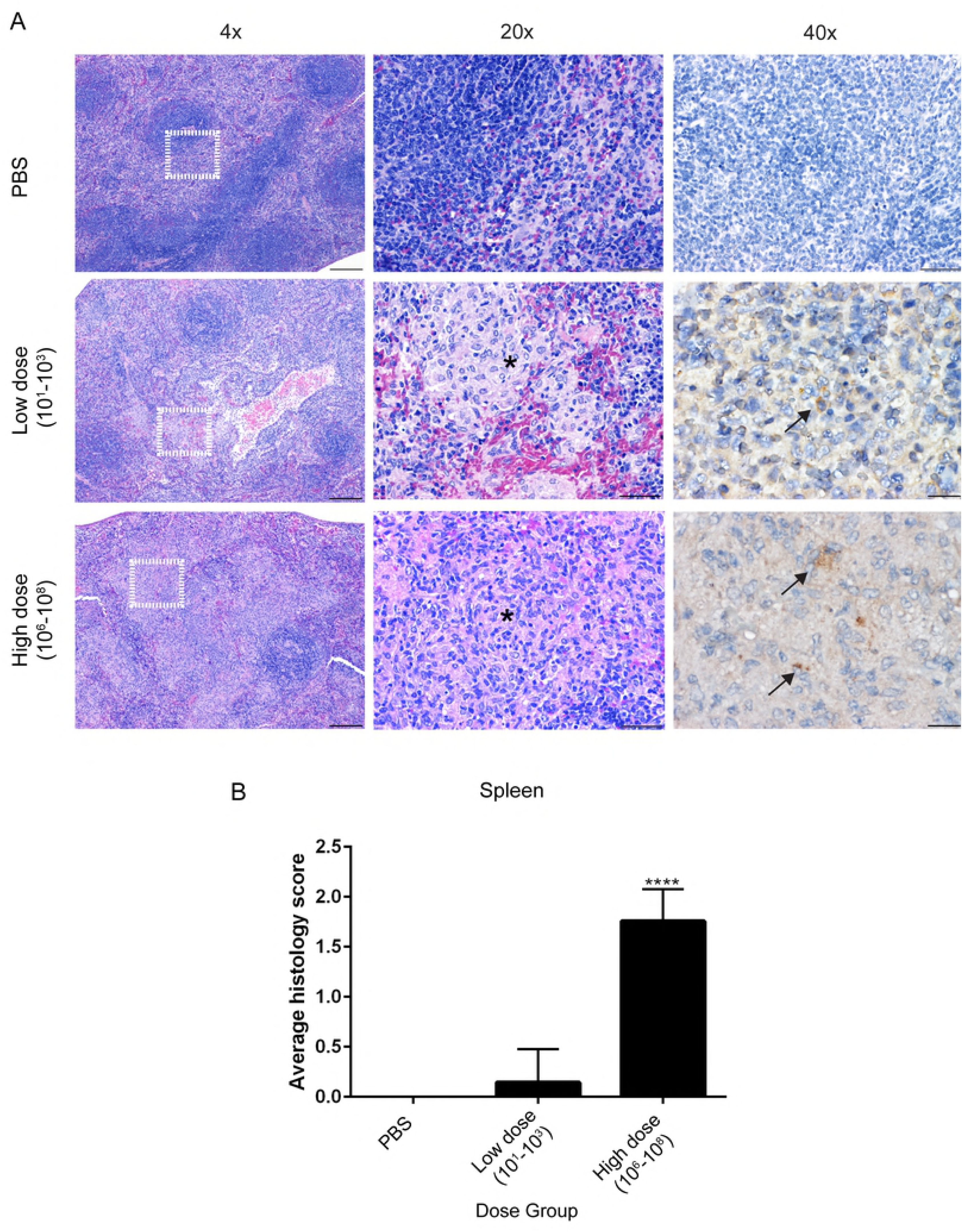
Spleen histopathology. Representative images of histopathology and immunohistochemistry of the spleen following intratracheal inoculation with PBS (top), *B. melitensis* 16M at low dose (middle), or high dose (bottom) at 30-days post-inoculation (A). Sections were scored for severity from 1-4 (Table S1) based on accumulation of epithelioid macrophages, neutrophils, and necrosis (B). Infection with *B. melitensis* induces accumulation of epithelioid macrophages (*). *Brucella* antigen was detected within epithelioid macrophages by immunohistochemistry (arrows). Magnification 4x (left, H&E, bar= 200 µm), 20x (middle, H&E, bar= 50 µm), 40x (right, Anti-*Brucella* IHC, bar=20 µm).

**Figure 6.**
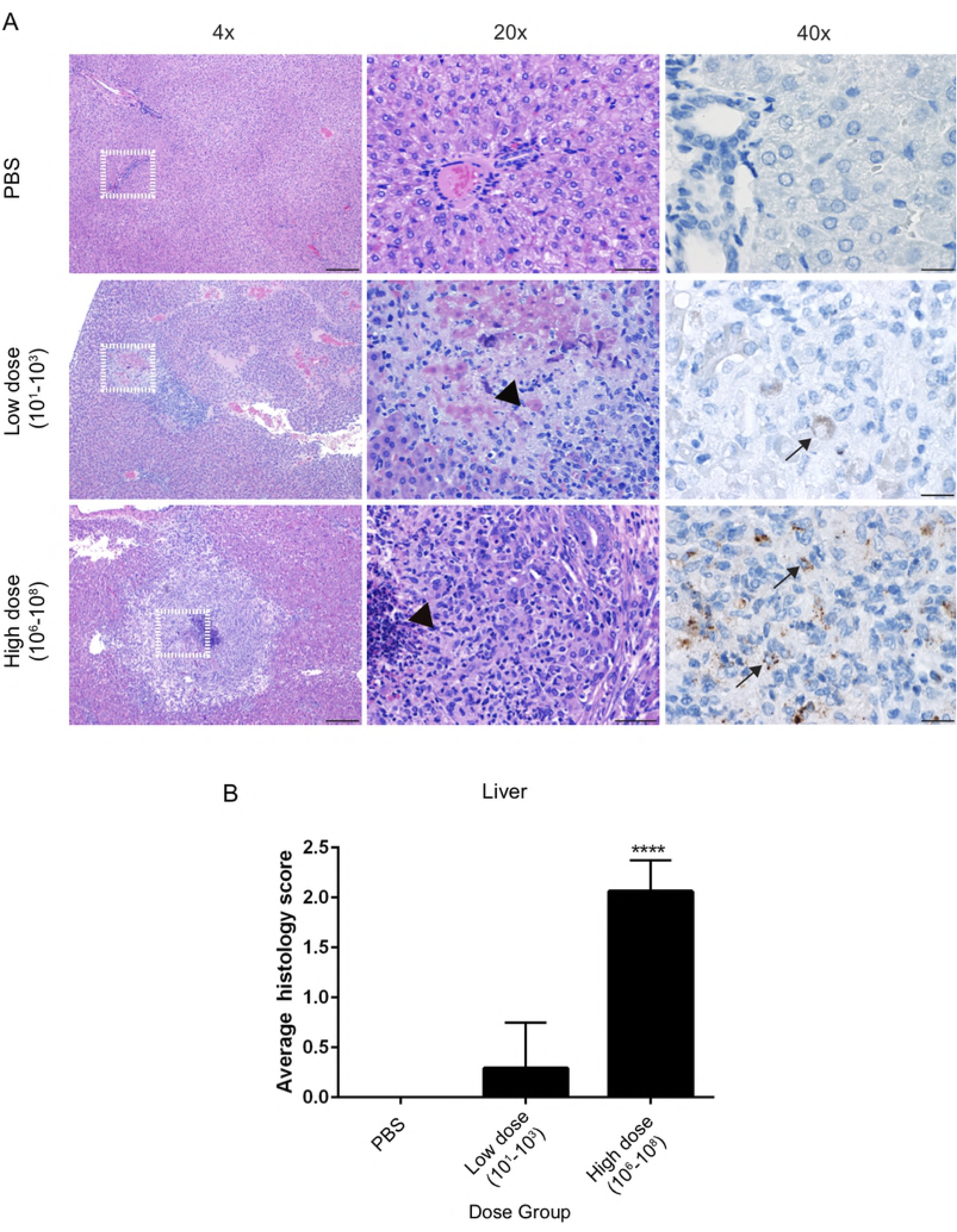
Liver histopathology. Representative images of histopathology and immunohistochemistry of the liver following intratracheal inoculation with PBS (top), *B. melitensis* 16M at low dose (middle), high dose (bottom) at 30-days post-inoculation. Sections were scored for severity from 1-4 (Table S1) based periportal inflammation, number and size of microgranulomas and necrosis. Foci of necrosis were seen in the low and high dose groups (arrowheads), but the lesions were larger in the high dose group. *Brucella* antigen was detected within necrotic hepatocytes and macrophages in areas of necrosis by IHC (arrows). Magnification 4x (left, H&E, bar= 200 µm), 20x (middle, H&E, bar= 50 µm), 40x (right, Anti-*Brucella* IHC, bar=20 µm).

*Brucella* species have a known tropism for the reproductive tract. In the natural host (sheep and goats), *B. melitensis* causes midterm spontaneous abortion and fetal death [10]. Lesions of the non-pregnant uterus have not been reported in the guinea pig model previously, and most reproductive studies have focused on pregnant animals. Interestingly, at 2-weeks post-inoculation, the endometrial stroma was variably expanded by edema, and endometrial glands were distended by an inflammatory infiltrate of intact and degenerate neutrophils and macrophages. The lesion progressed in severity and by 3 and 4-weeks post-inoculation, foci of histiocytic inflammation were developing in the myometrium (Fig. 7). A single animal in the 10^8^-dose group had histiocytic salpingitis. No lesions were identified 1-week post-inoculation in the uterus.

**Figure 7.**
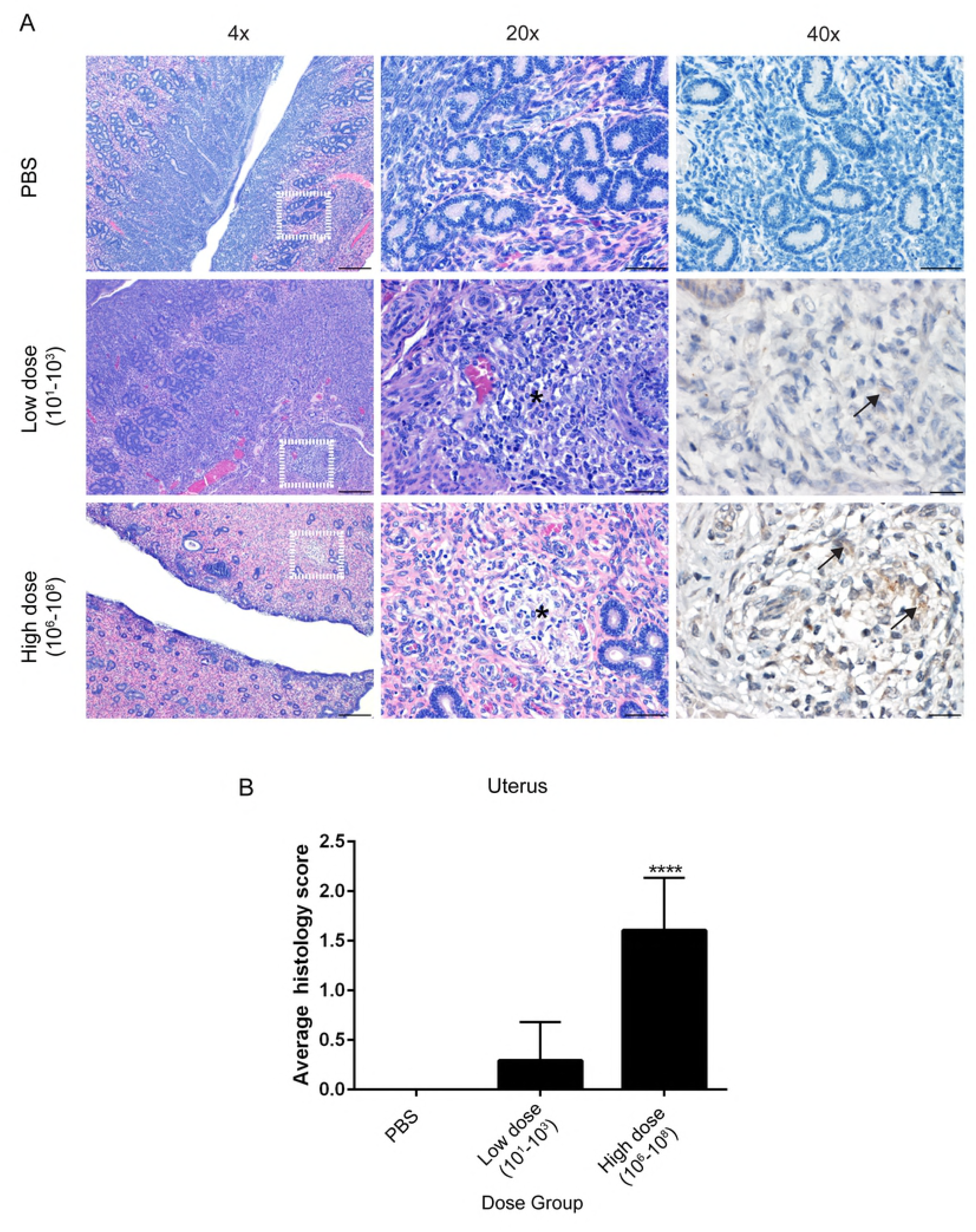
Uterine histopathology. Representative images of histopathology and immunohistochemistry of the uterus following intratracheal inoculation with PBS (top), *B. melitensis* 16M at low dose (middle), or high dose (bottom) at 30-days post-inoculation. Sections were scored for severity from 1-4 (Table S1) based on edema, endometrial neutrophilic inflammation, and myometrial inflammation. The high dose group had increased numbers of neutrophils in the endometrium, foci of histiocytic inflammation within the myometrium (*), and *Brucella* antigen was detected intracellularly via IHC (arrows). Magnification 4x (left, H&E, bar= 200 µm), 20x (middle, H&E, bar= 50 µm), 40x (right, Anti-*Brucella* IHC, bar=20 µm).

To further support the CFU data that the lesions in the liver, spleen, and uterus were due to *Brucella* infection, IHC was performed to colocalize *Brucella* antigen within foci of inflammation. *Brucella* antigen was detected within epithelioid macrophages in the spleen, liver, and uterus by IHC further corroborating the etiology of the lesion (Fig. 5-7). Antigen was also detected intracellularly within macrophages in the lung (Fig. S2), CLN, and TBLN (data not shown).

### Infection stimulates a *Brucella*-specific IgG humoral immune response

Guinea pigs develop a humoral response (anti-*Brucella* specific IgG) to infection with *Brucella melitensis* delivered via intratracheal inoculation. No change in IgG level was noted in the PBS, 10^1^, or 10^2^ groups. Only guinea pigs in the high dose groups (10^6^, 10^7^, 10^8^) were capable of mounting a humoral response against *B. melitensis*. The increase in IgG level was statistically significant in the 10^7^ and 10^8^ groups at 4-weeks post-inoculation (*P* < 0.01) (Fig 8). Levels of *Brucella*-specific IgG antibodies increased starting 1-week post-inoculation and increased throughout the study period (data not shown).

**Figure 8.**
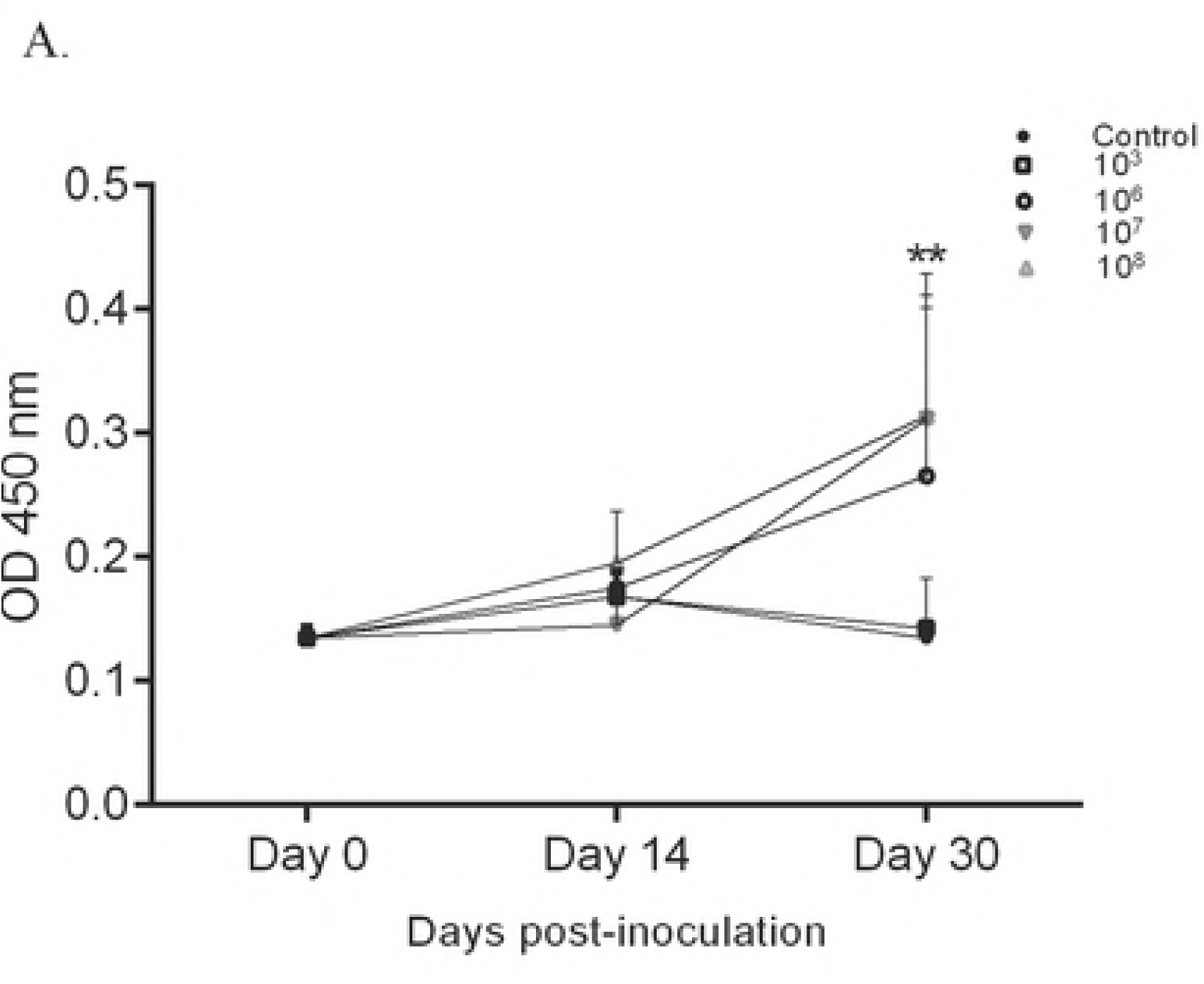
Humoral response to intratracheal inoculation with *B. melitensis*. Anti-*Brucella* specific IgG ELISA with sera from guinea pigs inoculated by intratracheal route with *B. melitensis* 16M at doses of 10^3^, 10^6^, 10^7^, 10^8^, or uninfected control (A) at day 0, 14, and 30 post-inoculation. Guinea pigs in the 10^7^ and 10^8^ groups developed a statistically significant humoral response to inoculation with *B. melitensis*. The results are expressed as the mean absorbance per group (± standard error). Statistical significance was determined by ANOVA followed by Dunnett’s multiple-comparison of each group (n=4) to the uninfected controls (n=3). Two asterisks, *P* < 0.01.

## Discussion

*Brucella* organisms can be easily aerosolized and inhalation of bacteria is a major route of natural transmission in both animals and people [4]. One of the limitations to developing stronger intervention measures such as a safe and efficacious vaccine has been the difficulty of replicating natural disease in a laboratory animal model. Mice are the most commonly utilized animal, but limitations to this model include lack of fever response, relatively high dose required to generate systemic infection, and an artificial route of inoculation that does not mimic natural transmission events [6]. In contrast, guinea pigs develop key features of disease when inoculated via an aerosol route, which closely mimics the naturally occurring disease process [13, 16, 17]. Guinea pigs were used in the early twentieth century as the model of choice to evaluate the pathogenicity of *Brucella* species such as *B. abortus*, *B. suis*, and *B. melitensis* and could offer an improvement over the mouse model for vaccine and therapeutic development [10, 11, 13, 16-24].

The PennCentury^TM^ MicroSprayer is a targeted means of generating aerosols and offers an improvement over aerosol chambers or aerosol devices like the Henderson apparatus because it allows for the direct inoculation of bacteria into the upper respiratory tract through the trachea. However, the MicroSprayer^®^ Aerosolizer does bypass the nares, which would be a line of defense in the upper respiratory tract against natural transmission. Due to the guinea pig oral anatomy, the device is inserted into the proximal trachea at the level of the arytenoid cartilage. Microparticles are generated after passage through the MicroSprayer^®^, which then move by centripetal force through the trachea and into the lower airways. This is similar to the natural transmission in which inhaled particles must pass from the nares into the trachea and then into the bronchi and bronchioles. Particle size determines the site of deposition within the airway with larger particles (>15 µm) removed through the nares and sinuses while smaller particles (6-10 µm) deposit in the bronchi [25]. The smallest particle size (≤5 µm) are able to deposit in the terminal bronchioles and alveoli [25]. The MicroSprayer^®^ generates a mean particle size of 8 µm, which allows for the particles to be deposited in the lower airways [26].

Recurrent or undulant fever is a hallmark of brucellosis in humans and is a feature of disease that is not replicated in the mouse model [2]. The first study to document fever in guinea pigs used an intraperitoneal, intravenous, or subcutaneous route of inoculation. The severity of the temperature elevation was not reported, and it was further stated that fever developed in the acute stage of infection, described as 72 hours post-inoculation [11]. In people, the onset of clinical symptoms such as fever tend to be insidious but likely develop between 6 to 90 days after exposure, and the temporality and undulant nature of the fever response suggests guinea pigs could be a biologically relevant model for future studies [4]. The aerosol literature with *Brucella* spp. in guinea pigs did not evaluate body temperature and thus it was previously unknown if aerosol inoculation would result in fever.

People can be infected with as few as 10-100 CFU of *Brucella* and thus this study evaluated the ability of low doses (10^1^, 10^2^,10^3^) of *B. melitensis* 16M to infect guinea pigs [4]. A high dose range (10^6^, 10^7^, 10^8^) was also evaluated because many of the infectious aerosols that people are exposed to likely exceed the minimum dose estimated to generate infection [2, 4]. The dose titration study indicated a dose of at least 10^6^ CFU was required to induce temperature elevations although systemic infection developed in the majority of the guinea pigs inoculated with 10^3^ CFU. Previous aerosol studies in guinea pigs delivered a dose of between 4.5×10^3^/ml to 5.0×10^5^/ml, which generated an estimated dose range of 48-2800 CFU [13, 16, 17, 27]. The majority of the early aerosol studies utilized the Henderson apparatus for generating aerosols, which is a mask that fits over the head and neck of the guinea pig to create a small aerosol chamber [13, 16, 17, 28]. As such, the guinea pigs were exposed not only through the respiratory tract, but bacteria were also likely deposited on mucous membranes of the conjunctiva and oral cavity and potentially ingested. The calculated dose did not account for these other potential routes of exposure, which could have increased the dose inoculated. Furthermore, since the doses from the earlier aerosol studies also based inoculation dose on calculations of ventilation rate and respiratory tidal volume of the guinea pig, the dose could have been underestimated [29]. The dose in this study is higher than the reported range required to induce infection in guinea pigs because we wanted to establish a model that replicated the features of human brucellosis like fever. The previous studies evaluated infection by colonization of organs such as the spleen and liver, whereas this study used clinical parameters such as body temperature plus organ colonization to demonstrate infection.

While the respiratory tract is a common portal of entry, pulmonary pathology and respiratory disease are not atypical features of *Brucella* spp. infection [30, 31]. In the rare cases in which respiratory disease is reported, the common presentations include pneumonia, bronchopneumonia, pleural effusion, and dry coughing [31]. Respiratory signs rarely occur in isolation, and patients often have concomitant disease such as hepatitis or spondylitis supporting the role of the lung as a portal of entry rather than a primary target [30, 31]. Clinical signs in mice with respiratory infection have not been reported [7]. In the 10^8^ group, two animals developed transient ocular discharge, which can be associated with respiratory disease in guinea pigs [32]. Guinea pigs had a pattern of cranioventral lung lobe consolidation and embolic foci, which suggests a dual pattern of infection. The initial inoculation with *B. melitensis* via intratracheal delivery likely leads to the development of cranioventral consolidation as the site of initial deposition followed by an embolic pattern as the animals become bacteremic. A previous aerosol study in guinea pigs by Elberg and Henderson reported no *Brucella*-specific macroscopic or microscopic pulmonary pathology, and several other contemporary studies failed to evaluate the lung for lesions [13, 16, 17].

*Brucella* has a tropism for organs of the reticuloendothelial system including the spleen, lymph nodes, and liver [2, 3, 14, 15]. Splenomegaly, lymphadenomegaly, and hepatitis are common macroscopic lesions in natural and experimental infection [3]. The microscopic splenic lesion has not been well described in the medical literature but has been described as congestion, lymphoid hyperplasia, and histiocytic splenitis in mice [6]. Guinea pigs develop splenic congestion and lymphoid hyperplasia with occasional necrosis and abscesses thirty days after receiving an aerosol dose of 2.16×10^3^ CFU of *B. abortus* and *B. melitensis* [13].

Lymphadenomegaly is another well documented sequelae of infection with *Brucella* spp. in both people and guinea pigs [13, 15-17]. An aerosol study by Elberg and Henderson noted the development of caseous abscesses in the cervical and tracheobronchial lymph nodes; however, *Brucella* was not cultured from the nodes so the etiology of the abscess cannot be definitively assigned to brucellosis [13]. The final reticuloendothelial organ that is commonly affected during infection is the liver. A prospective study of patients with hepatitis due to *B. melitensis* found that disease is often subclinical but can cause mild derangements in hepatic enzymes such as alanine aminotransferase (ALT)[14]. The acute lesion of brucellosis is described most frequently as lymphocytic portal to lobular inflammation with fewer cases diagnosed with noncaseating granulomas or microgranulomas [14]. The literature describes “granulomata” in the guinea pig liver following aerosol inoculation but do not provide further histologic description [13].

*Brucella* spp. are best known as pathogens of the reproductive tract during pregnancy and cause a range of adverse events such as abortion, stillbirths, and infertility in small ruminants and people [2, 33]. Less is known about the tropism of *Brucella* organisms for the non-gravid uterus. Reproductive studies in mouse models have not reported lesions in non-pregnant female reproductive organs. Researchers in the early twentieth century did not identify lesions in the reproductive tract of female guinea pigs and thus it was assumed that females were not an appropriate animal model for use in reproductive pathogenesis investigations. Instead, the early studies focused on male guinea pigs and identified orchitis, epididymitis, and peri-orchitis subsequent to intraperitoneal, intratesticular, and aerosol inoculation [10, 11, 22, 24, 34]. However, a study from 1974 demonstrated that when pregnant guinea pigs are inoculated at mid-gestation with 10^5^ *B. abortus* 2308 via intramuscular injection, stillbirths, abortions, and vertical transmission occur. Thus, guinea pigs may be suitable models for future investigations into the pathogenesis and tropism of *Brucella* spp. for the gravid uterus.

This study describes pathologic changes of the non-gravid uterus, broadens the knowledge of *Brucella* as a pathogen of the reproductive tract, and suggests that pregnancy is not required to generate tropism. Since infertility is also described in non-pregnant women infected with *Brucella* spp., it is possible that inflammation of the reproductive tract is a contributing factor [33]. Intratracheal inoculation of the guinea pig offers an intriguing model for the study of the host-pathogen interaction that leads to reproductive disease in addition to providing a reliable means of generating systemic and clinical brucellosis that could be used to evaluate vaccine candidates.

## Methods

### Ethics statement

This study includes the use of guinea pigs. This study was carried out in an approved facility in strict accordance with all university and federal regulations. All guinea pig experimentation was reviewed and approved by the Texas A&M University Laboratory Animal Care and Use Committee (protocol: 2015-0036). The protocol was approved and is in accordance with the Institutional Animal Care and Use Committee (IACUC) policies of Texas A&M University. Texas A&M is accredited by the Association for the Assessment and Accreditation of Laboratory Animal Care, International (AAALAC).

### Animal husbandry

Outbred Harley female guinea pigs (n=44) weighing approximately 300-350 g were obtained from Charles River Laboratories and housed in microisolator caging in a biosafety level three facility. Guinea pigs were acclimated to the facility for 5 days prior to infection and were on a 12-hour—12-hour light-dark cycle with ad libitum access to pelleted food, Timothy hay, and water. A modified Karnofsky performance status scoring system was used to evaluate the guinea pigs daily to determine if early removal from the study was required.

### Bacteriology

*Brucella melitensis* 16M wild-type strain, originally acquired from an aborted goat fetus, was routinely grown on tryptic soy agar (TSA) (Difco Laboratories) at 37°C in an atmosphere containing 5% (vol/vol) CO2 for 72 hours [35]. Bacteria were harvested into phosphate-buffered saline (PBS) (pH 7.4; Gibco) to obtain the final concentration needed for each experiment, as estimated turbidometrically using a Klett meter. Serial dilution was performed to accurately determine the number of organisms in the inoculum. To determine if passage through the MicroSprayer^®^ affected the inoculum dose, 100 µl of the inoculum was passed through the MicroSprayer^®^ and collected in the microcentrifuge tube containing 900 µl PBS for serial dilution and culture on TSA.

### Dose titration

Guinea pigs were divided in 7 groups (n=4) and were further subdivided into low dose (10^1^, 10^2^,10^3^), high dose (10^6^, 10^7^, 10^8^), or control (PBS) groups. Guinea pigs were anesthetized with ketamine/xylazine (50mg/kg;5mg/kg) and a subcutaneous IPTT-300 microchip was placed to monitor temperature throughout the study (Bio Medic Data Systems). The 50 µl doses of *B. melitensis* 16M were prepared from cultures resuspended into PBS and serially diluted to obtain the dose groups. The inoculum was administered into the proximal trachea and lungs using the PennCentury^TM^ MicroSprayer I-1C device (Penn Century Inc.). Animals were monitored daily for 30 days for changes in body temperature, respiratory pattern and effort, and weight. Temperatures of ≥39.5°C were defined as fever. At 30-days post-inoculation, animals were euthanized by intraperitoneal injection of sodium pentobarbitol (Beuthanasia) followed by cardiac exsanguination. Samples of lung, liver, spleen, cervical lymph node, tracheobronchial lymph node, and uterus were aseptically collected into 1 ml PBS, homogenized, serially diluted, and 100 µl of each dilution was plated in duplicate onto Farrell’s medium (TSA plus *Brucella* Oxoid supplement, equine serum, and 50% dextrose) and incubated at 37°C in an atmosphere containing 5% (vol/vol) CO2 [7]. Bacterial colonies were enumerated after 72 hours to quantify tissue colonization. Spleen and liver were weighed at necropsy, and the aforementioned tissues were collected and fixed in 10% neutral buffered formalin for evaluation by light microscopy.

### Kinetics of infection

Guinea pigs were divided into four groups (n=4) and were infected via intratracheal inoculation with 50 µl of 1×10^7^ CFU *B. melitensis*. The endpoints were 2-hours post-inoculation and at weekly intervals thereafter for three weeks. To determine the actual number of infectious organisms delivered by intratracheal inoculation, 4 animals were euthanized 2-hours post-inoculation, and the lung was divided into four quarters (left and right, cranial and caudal), collected into 1 ml PBS, homogenized, and serial dilutions plated on Farrell’s medium. Spleen, liver, CLN, TBLN, and uterus were collected for culture and histology at each of the time points, as described in experiment 1.

### Histopathology

Spleen, liver, lung, uterus, CLN, and TBLN were collected at necropsy and fixed in 10% neutral buffered formalin for a minimum of 48 h. Tissues were routinely processed and embedded, sectioned at 5 µm, and stained with hematoxylin and eosin. Sections from spleen, liver, lung, and uterus were graded in a blinded fashion by a board-certified veterinary pathologist (MH) on a scale of 0-4 for inflammation type, necrosis, and severity (S1). The mean total score for each tissue was compared between groups.

### Immunohistochemistry

Unstained slides from spleen, uterus, liver, and lung were adhered to positively charged glass slides for immunohistochemistry. Slides were deparaffinized and rehydrated through a series of xylene and ethanol steps before antigen retrieval was performed using 1:10 EMS Solution A (Electron Microscopy Services) in a 2100 Antigen Retriever (Aptum Biologics Ltd.), according to manufacturer protocol. Endogenous peroxidases were blocked by 10 m incubation with Bloxall Blocking Solution (Vector Laboratories) followed by 20 m blocking with normal goat serum (Vector). After each step slides were washed with PBS plus 0.5% tween for 5 minutes. Primary incubation was overnight at 4°C with *Brucella* polyclonal rabbit antibody (Bioss) at 1:600. Negative control tissues were incubated with rabbit nonimmune serum diluted in PBS. A Vectastain ABC and Betazoid DAB chromagen kits (Biocare Medical) were used following primary incubation according to the manufacturer’s instructions. The slides were counterstained with Meyer’s hematoxylin III.

### Anti-*Brucella* specific IgG ELISA

300 µl of blood was collected into serum separator tubes from the lateral saphenous vein at day 14 and from the heart at day 28 following euthanasia. Blood was centrifuged at 3000 rpm for 5 minutes, and the serum was collected for anti-*Brucella* specific immunoglobulin G (IgG) indirect enzyme linked immunosorbent assay (iELISA). 96 well plates were pre-coated with 25 µg/well of *Brucella abortus* 2308 heat killed lysate and held overnight at 4°C. Plates were washed three times and then blocked with 3% skim milk (Sigma) for 2 hours at room temperature. Guinea pig sera samples were diluted in blocking buffer (0.25% [wt/vol] bovine serum albumin) to 1:1000 and incubated at 37°C for 1 h. Plates were washed five times and then peroxidase labeled goat anti-guinea pig IgG (KPL) was added at 1:2000, followed by incubation at 37°C for 1 hour. After a final washing step, horseradish peroxidase substrate (Sigma) was added and plates were protected from light and incubated for 30 m at 37°C. Absorbance was measured at 450 nm. All assays were performed in triplicate, and the results are presented as the mean value for the three wells.

### Statistical analysis

Analysis was performed using the GraphPad Prism 6.0 Software. The difference between group means was analyzed using a one-way analysis of variance (ANOVA) repeated-measures test, and Dunnett’s multiple comparisons was used to generate *P* values for selected mean comparisons. Tukey’s multiple comparison was used to generate *P* values to compare mean IgG values.

## Acknowledgements

We thank Anthony Gregory and Erin J. van Schaik for technical guidance and support.

## Supporting Information Legends

**Supplemental Table 1 Grading criteria for histopathology**

Spleen, liver, uterus, and lung were graded according to inflammation and lesion severity.

**S1 Fig Distribution of aerosol particles following intratracheal inoculation.**

Intratracheal inoculation results in even distribution of aerosolized particles throughout all lung fields. The distribution of aerosolized *B. melitensis* 16M in the lung lobes of guinea pigs inoculated with 1×10^7^ CFU/50 μl was evaluated at 2-hours and 1,2, and 3-weeks post-inoculation. The lung was divided into four regions defined as left, right, cranial, and caudal, and tissue colonization was determined by region. The horizontal bar is the mean per group with standard error.

**S2 Fig Pulmonary histopathology.**

Representative images of histopathology and immunohistochemistry of the lung following intratracheal inoculation with PBS (top), *B. melitensis* 16M at low dose (middle), high dose (bottom) at 30-days post-inoculation. Sections were scored for severity from 1-4 (Table S1) based neutrophilic inflammation, number and size of microgranulomas and necrosis, and bronchoalveolar hyperplasia. Foci of histiocytic inflammation were seen in the low and high dose groups (arrowheads), but the lesions were larger in the high dose group. *Brucella* antigen was detected within alveolar macrophages in areas of inflammation by IHC (arrows). Magnification 4x (left, H&E, bar= 200 µm), 20x (middle, H&E, bar= 50 µm), 40x (right, Anti-*Brucella* IHC, bar=20 µm).

